# Temporal degradation of PRC2 uncovers specific developmental dependencies

**DOI:** 10.64898/2026.04.18.719361

**Authors:** Ming-Kang Lee, Sebastian D. Mackowiak, Daniel Felismino, Jeron Venhuizen, Maria Walther, Alexander Meissner

## Abstract

The repression of genes by Polycomb repressive complex 2 (PRC2) is well documented and constitutive deletion studies have highlighted its essential role in development. However, exactly how loss of the complex is linked to these phenotypes remains incompletely understood and difficult to study in vivo. Here, we applied a rapid protein degradation strategy combined with a scalable embryoid model to trigger temporal PRC2 loss. This allowed us to fine-map developmental failures and showed that in addition to the canonical posterior phenotypes, PRC2-depleted embryoids display unexpected ectopic expression of anterior and lateral lineage genes. We also find that while the baseline enrichment of H3K27me3 generally predicts lineage sensitivity, the presence of cognate transcription factors ultimately helps explain which genes respond to PRC2 loss. Our integrative expression and chromatin profiling highlights that in contrast to developmental genes, pluripotency-associated genes only fail to silence with early PRC2 depletion but are unaffected once cells exited pluripotency. Together, our results shed new light on the specific developmental requirements for PRC2-mediated repression at unprecedented temporal and lineage resolution.

## Main

The coordination of lineage-specific transcription factors (TFs) and ubiquitously expressed epigenetic regulators orchestrates development and maintains cellular identities^1^. In mammals, depletion of chromatin modifying enzymes is often lethal around gastrulation^2–4^, which emphasizes their essential role as the complexity of the developing embryo increases. This includes the Polycomb Repressive Complex 2 (PRC2), which contains four core subunits (Ezh1/2, Suz12, Eed, RbAp46/48) and mediates gene silencing via histone H3 lysine 27 trimethylation (H3K27me3)^5–8^. In pluripotent cells, the combination of H3K27me3 and H3K4 trimethylation (H3K4me3) establishes a “bivalent” state at promoters of most developmental genes, which is thought to keep genes in a poised state for activation upon receiving differentiation cues^9–12^. Loss-of-function mutations of PRC2 core subunits in pluripotent stem cells cause general de-repression of bivalent genes, and constitutive disruption *in vivo* transforms the body plan posteriorly prior to embryo lethality^13–15^. However, the early inactivation in either germline or zygotic deletion studies makes it difficult to properly disentangle the impact of acute PRC2 loss as development proceeds.

### Temporal PRC2 degradation reveals stage-specific differentiation defects

To better understand PRC2s role in gastrulation, we employed our previously developed mouse embryonic stem cell (mESC)-based gastruloid model, termed trunk-like structures (TLSs)^16^. TLSs sufficiently recapitulate relevant morphological and spatiotemporal programs of post-occipital gastrulating embryos, exhibiting highly organized compartments, including a central neural tube, bilateral somites, endoderm, and bipotent neuromesodermal progenitors (NMPs) at the posterior pole that continuously give rise to neural and mesodermal tissues^17–19^.

As a reference to previous constitutive PRC2 disruptions, we used *Eed*-knockout (Eed-KO) mESCs to derive TLSs, which expectedly fail to develop and show massive cell death (**Extended Data Fig. 1a-c**). To acutely inactivate PRC2 activity, we integrated the FKBP-based degradation tag (dTag) into the *Suz12* locus of dual reporter mESCs with a T::H2B-mcherry/ Sox2::H2B-Venus, which facilitate image-based monitoring of mesodermal and neural lineage composition following PRC2 loss^16,20,21^. Our western blot analysis confirms that efficient degradation of SUZ12 occurs already 2 hours after treatment with degrader dTag^V-1^ and H3K27me3 levels become nearly undetectable after 24 hours of treatment (**Fig. 1a**). As a negative control, we treated cells with its diastereomer (dNeg), which had no effect on PRC2 or H3K27me3. We also confirmed the similar depletion of SUZ12 and H3K27me3 in 3D structures after four days of TLS development and a 24-hour treatment window (**Fig. 1b, Extended Data Fig. 1d**).

**Fig. 1:**
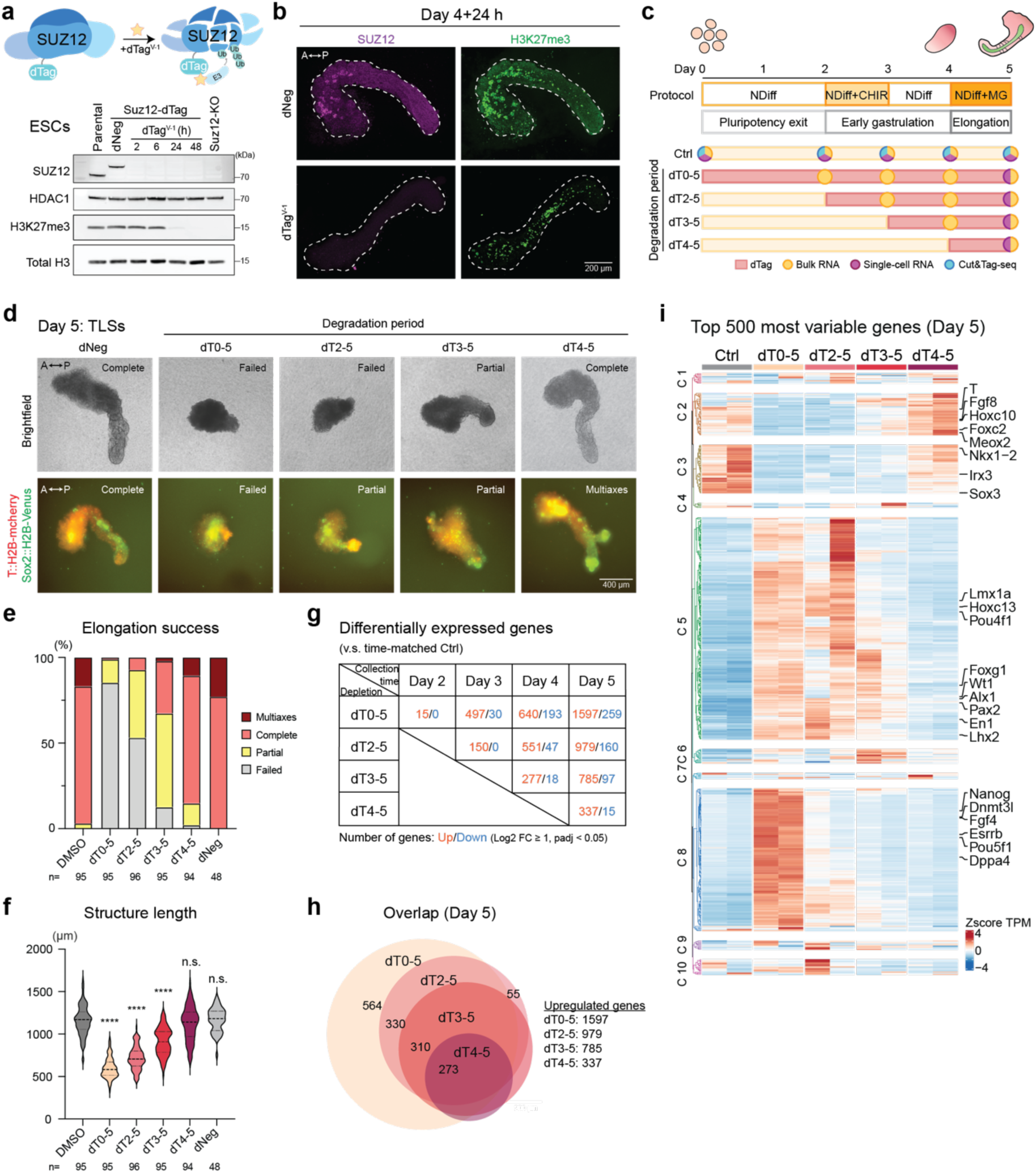
Acute PRC2 depletion causes activation of ectopic lineage genes. a, Schematic of Suz12-dTag in mouse ES cells. Western blot showing depletion of SUZ12 and H3K27me3. HDAC1 and total H were used as loading controls **b,** Maximum intensity projection of z-stacked whole-mount immunofluorescence showing SUZ12 degradation in the TLS system. **c,** Schematic of SUZ12 depletion conditions during TLSs generation. Red color bars denote SUZ12 depletion. Dots denote sample collection. NDiff, N2 and B-27 neural differentiation media; CHIR, WNT agonist CHIR99021; MG, Matrigel. **d,** Live imaging of TLSs under depletion conditions. A, anterior; P, posterior. **e,** Bar plot showing the percentage of elongation success. The structures were collected from 2 independent experiments. **f,** Violin plot representation of structure length across depletion conditions. The median is indicated by a dashed line, and the interquartile range is demarcated by dotted lines. **g,** Table showing numbers of differentially expressed genes comparing PRC2-depleted TLSs to time-matched controls. Differential analysis is based on a threshold of Log2 fold change (Log2 FC) ≥ 1 and *P*_adj_ < 0.05. **h,** Venn diagram displaying the overlap of upregulated genes across depletion conditions in TLSs at 5 dpa. **i,** Hierarchical clustering heatmap showing the top 500 most variable genes in TLSs at 5 dpa. Clustering was performed using Euclidean distance metrics. Color intensity represents Z-score normalized TPM levels.

To take full advantage of our model, we next set up multiple starting points for dTag treatment to perturb PRC2 activity at specific timepoints during gastrulation. These include depletion initiated at the time of cell aggregation (dT0-5; treatment starting at T0 and proceeding continuously until day 5), or at two (dT2-5), three (dT3-5), and four days (dT4-5) post-aggregation (dpa) (**Fig. 1c**), followed by morphological, bulk and single-cell transcriptomic as well as epigenetic analyses. Regarding stage selection, the pulse of WNT agonist CHIR99021 between 2-3 dpa is thought to induce a primitive streak (PS)-like stage, promoting differentiation of pluripotent cells into defined germ layers (**Fig. 1c**)^22^. Symmetry breaking due to the protrusion of an elongating front can be observed by the end of 4 dpa (**Extended Data Fig. 1e**)^16^. Lastly, the addition of Matrigel on day 4 leads to formation of a central neural tube, bilateral somites, and Sox2/T double-positive NMPs at the posterior end (**Fig. 1d**). Under the early-onset PRC2 depletion (dT0-5 and dT2-5), the majority of the structures exhibit delayed symmetry breaking and eventually show failed or only partial elongation (**Fig. 1d-f, Extended Data Fig. 1e-f**). Although the later PRC2 depletion (dT3-5) does not delay symmetry breaking, most structures still display defective elongation with reduced structure length compared to the dNeg treatment. The stage around the activation of WNT signaling appears especially sensitive to the loss of PRC2 and generally exhibits more severe morphological defects including lack of elongation. The observed phenotypic defects were accompanied by cell death that seems to more prominently affect mesodermal cell types (**Extended Data Fig. 1g**).

Our findings are in line with prior work showing that undifferentiated PRC2-deficient mESCs were viable, but showed compromised embryoid body and neural progenitor cell (NPC) differentiation^23,24^. However, through the temporal depletion, we could uncover specific periods of enhanced sensitivity and more nuanced lineage-specific effects.

### PRC2 loss upregulates ectopic lineage genes

To determine how PRC2 loss disrupts developmental programs, we performed time-resolved bulk RNA-sequencing (RNA-seq) and reconstructed the temporal transcriptional dynamics across depletion conditions. The transcriptomic profiles of our control TLSs (treated with dNeg) are reminiscent of those of gastrulating embryos (**Extended Data Fig. 2a-b**)^16,22^. PRC2-depleted TLSs initially follow similar differentiation trajectories to the control TLSs (**Extended Data Fig. 2c**). However, we find that the duration of PRC2 depletion correlates with increasing levels of dysregulation, leading to more, largely overlapping, differentially upregulated genes (**Fig. 1g-h, Extended Data Fig. 2d, Supplementary Table 1**). Expectedly, only a minor fraction of genes is downregulated, aligning with PRC2’s role as a transcriptional repressor (**Extended Data Fig. 2e**).

One classic hallmark of Polycomb mutant embryos is the disruption of the spatiotemporal *Hox* collinearity, an important determinant of body plan formation. Posterior *Hox* genes become ectopically expressed in the more anterior segments, causing a “posterior homeotic transformation”^8,25–29^. To test if we observe a similar phenomenon, we compared the temporal expression pattern of the *Hoxa* locus (**Extended Data Fig. 2f**). Early depletion caused excessive expression across the *Hoxa* locus at 3 dpa. At the final stage of TLS, upregulation is confined to the posterior-most *Hoxa13*, while anterior *Hoxa* genes become downregulated with prolonged PRC2 depletion. Interestingly, PRC2 depletion post-WNT activation (dT3-5) does not induce any major *Hox* de-repression, but causes nonetheless substantial elongation defects.

To determine the degree of transcriptional dysregulation, we integrated the top 500 most variable genes at 5 dpa into a common heatmap and identified clusters of genes with common expression patterns (**Fig. 1i**). PRC2-depleted TLSs show reduced expression of NMP (*T, Fgf8, Nkx1-2*), somite (*Foxc2, Meox2*), and neural tube genes (*Irx3, Sox3*; Clusters 2 and 3), which is consistent with the compromised elongation. Gene Ontology (GO) analysis confirms that these genes are involved in anterior-posterior patterning, segmentation, and somitogenesis (**Extended Data Fig. 2g**). A specific cluster of genes (Cluster 8) is highly expressed exclusively under the earliest onset of PRC2 depletion (dT0-5). Many of these genes are associated with stem cell maintenance and pluripotency. Interestingly, once silenced from day 2 onwards, loss of PRC2 activity no longer induces their reactivation, indicating that the impact of PRC2 loss is transient and other epigenetic layers become subsequently more relevant. However, we inferred that the persistence of pluripotency is not the primary cause of gastrulation failure, as this was not observed in other depletion conditions. Another cluster of genes (Cluster 5) are consistently upregulated across all depletion conditions, suggesting an ongoing dependence on PRC2-mediated repression. This includes genes associated with epithelial tube morphogenesis (*Wt1, Pax2, Lhx1/2*), limb development (*Alx1/3/4, Shox2),* and brain development (*En1*, *Lmx1a, Pou4f1, Foxg1*). To date, the TLS system lacks encephalic tissues, intermediate mesoderm (IM), and lateral plate mesoderm (LPM)^16,19,30^. Therefore, it is surprising that PRC2 depletion leads to upregulation of IM/kidney genes (*Lhx1/2, Pax2, WT1*), and LPM/limb genes (*Alx1/3/4, Shox2*)^31,32^. Even more surprisingly, the de-repression of anterior neural genes challenges the canonical paradigm of posterior transformation in Polycomb-mutant embryos.

Together, we conclude that developmental defects caused by acute PRC2 loss result from de-repression of ectopic lineage genes, compromising the cellular identities.

### Transcriptional heterogeneity increases in the absence of PRC2

Given the misexpression of other lineage markers, we wondered if PRC2 loss induces differentiation into alternative cell fates. We performed single cell-RNA sequencing (scRNA-seq) of mESCs, normal TLSs (2, 3, 4, and 5 dpa), and PRC2-depleted TLSs (dT0-5, dT2-5, dT3-5, dT4-5 at 5 dpa) and reconstructed the developmental trajectory under normal and perturbed conditions (**Fig. 2a, Extended Data Fig. 3a-c**)^33,34^. Using previously published gastruloid and TLS scRNA-seq datasets as references, we annotate 18 cell clusters, including epiblasts, primitive streak, and NMPs, which bifurcate into somitic and neural lineages (**Extended Data. Fig. 3d-e**)^16^. Interestingly, we do not identify any cluster consisting exclusively of PRC2-depleted TLSs, suggesting no novel cell types are induced despite expressing brain, kidney, and limb-related genes (**Extended Data Fig. 3f**). However, PRC2 inactivation leads to a decreased representation of NMPs as well as nascent mesodermal lineages (presomitic mesoderm, somite-1/0), which may be linked to a compromised stem cell reservoir at the posterior end. The extent of this decrease positively correlates with the duration of degradation, in line with the severity of elongation defects, mesoderm-specific sensitivity, and reduced NMP gene expression observed in our other analyses (**Fig. 2b**).

**Fig. 2:**
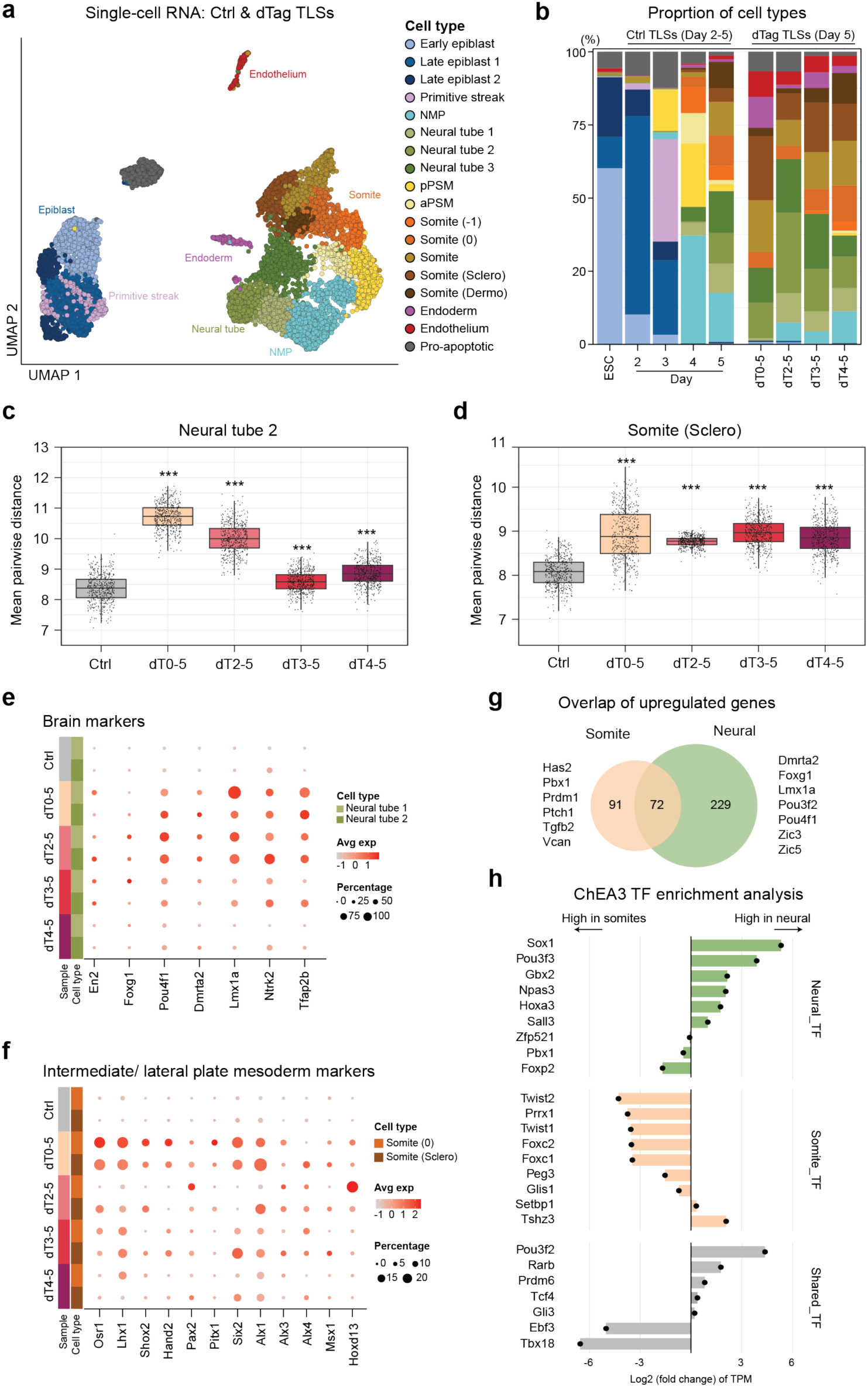
PRC2 loss leads to stochastic de-repression of lineage genes. **a**, Uniform manifold approximation and projection (UMAP) plot showing 18 cell types identified in control mESCs, TLSs at 2,3,4, and 5 dpa, and PRC2-depleted TLSs at 5 dpa. Each dot represents a cell. n= 9091 cells. **b,** Box plot representation of cell type proportion in normal TLSs across time points and PRC2-depleted TLSs at 5 dpa. The colors correspond to the cell types as represented in **a**. **c,** Box plot showing the mean pair-wise distance of 35 randomly selected cells within neural tube 2 in the UMAP plot. Each dot represents one round of iteration. Statistical significance was determined via one-way ANOVA. N = 500 iterations. **d,** Box plot showing the mean pair-wise distance of randomly selected cells within sclerotome, as in **c. e,** Dot plot displaying expression of brain markers in neural tube 1 and 2 across samples at 5 dpa. This size of the dots represents percentage of expressing cells and the color represents the expression level. **f,** Dot plot representation of the expression of IM and LPM markers in somite (0) and somite (sclerotome). **g,** Venn diagram representation of the overlap of upregulated genes in neural and somitic lineages, as analyzed in **Extended Data** Fig. 4d. **h**, Box plot displaying the Log2 fold changes in the expression of TFs in normal neural tubes and somites. The expression was represented by pseudobulk TPM in neural lineages (neural tube 1 and 2), and mesodermal lineages (somite 0, and somite sclerotome).

**Fig. 3:**
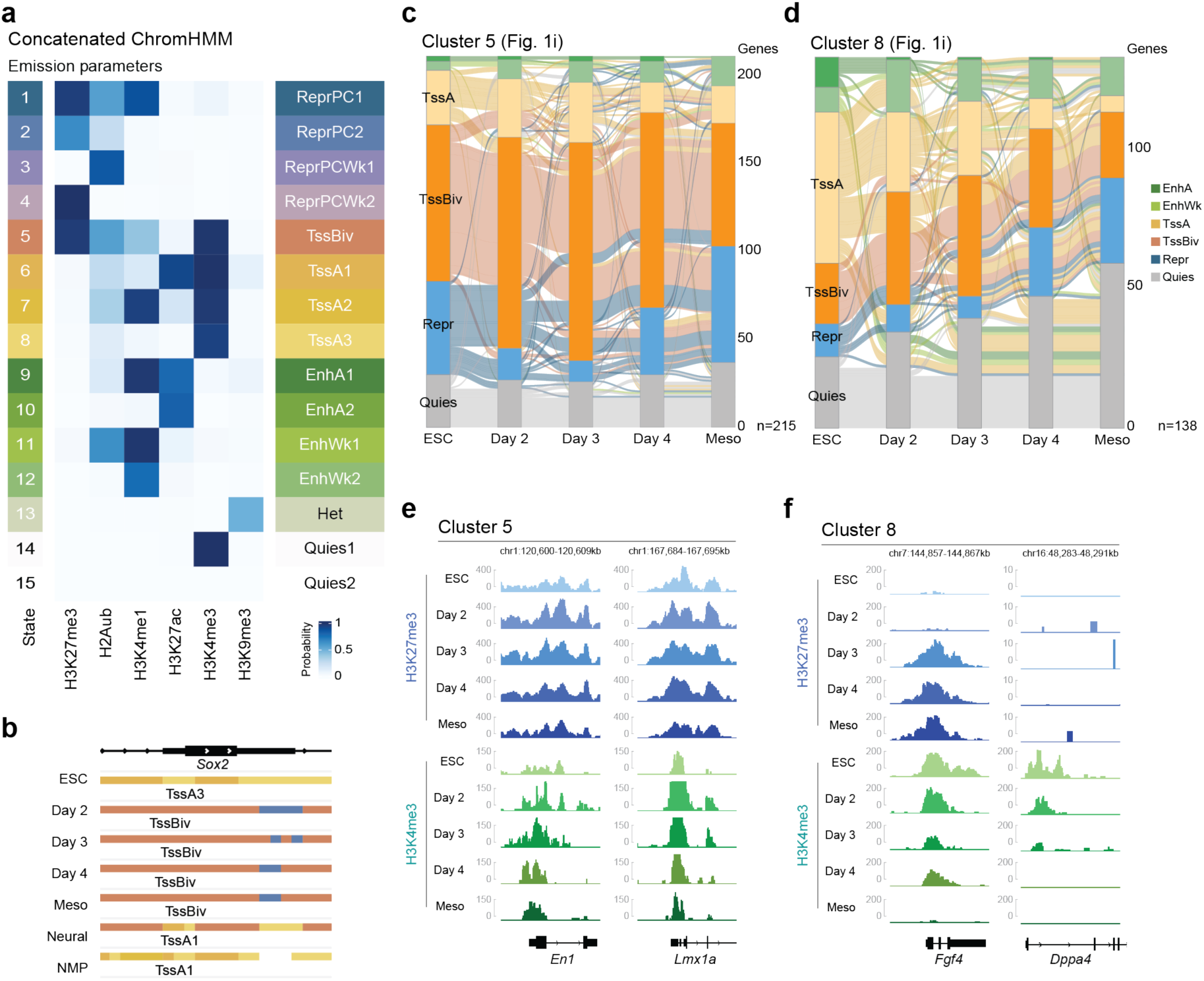
Chromatin state transitions define modes of upregulation. a, Heatmap displaying concatenated emission parameters across cell types. Each row corresponds to a different chromatin state, and each column corresponds to a different histone mark. The intensity of the color represents the probability of observing the mark in the state. ReprPC1, repressed by Polycomb 1; ReprPC2, repressed by Polycomb 2; ReprPCWk1, weakly repressed by Polycomb 1; ReprPCWk2, weakly repressed by Polycomb 2; TssBiv, bivalent transcription start site (TSS); TssA1, active TSS 1; TssA2, active TSS 2; TssA3, active TSS 3; EnhA1, active enhancer 1; EnhA2, active enhancer 2; EnhWk1, weak enhancer 1; EnhWk2, weak enhancer 2; Het, heterochromatin; Quies1, quiescence 1; Quies2, quiescence. **b**, Snapshot showing the chromatin states transition at the *Sox2* locus. **c,** Alluvial plot displaying the chromatin state transitions of Cluster 5 genes across time points. For visualization, similar states were collapsed into a single category, such as the four “repressed by Polycomb” states are represented as “Repr”, the three “active TSS” are represented by “TssA”. The colors represent the emission states as in **a**. **d,** Alluvial plot displaying the chromatin state transitions of Cluster 8 genes across time points. **e,** Genomic snapshots showing stable enrichment of H3K27me3 and H3K4me3 at *En1* and *Lmx1a* loci across samples. **f,** Genomic snapshots showing changes in H3K27me3 and H3K4me3 levels at the *Fgf4* and *Dppa4* loci across samples.

Next, we sought to explore transcriptional heterogeneity upon PRC2 loss and its possible effects on cellular identity^35^. We iteratively computed the mean pairwise distances of randomly selected cells within the same cell types (e.g., neural tube 2 and sclerotome) across depletion conditions^36^, and found that PRC2-depleted TLSs uniformly show greater distances (**Fig. 2c-d**). This prompted us to explore whether increased heterogeneity arises from the stochastic activation of ectopic lineage genes as observed in bulk RNA-seq. Selected brain marker genes are silent in the control neural tubes, but increase both in frequency and expression level in neural lineages upon PRC2 degradation, suggesting co-expression of ectopic marker genes from distinct but related lineages (**Fig. 2e**). A similar de-repression of IM and LPM genes is also observed in somitic tissues (**Fig. 2f**). The magnitude of the increase positively correlates with the depletion duration as expected. To ensure this is not an artifact of the *in vitro* system, we also analyzed previously published *in vivo* mouse *Eed*-KO and WT embryos scRNA-seq datasets^4^. Due to the lack of neural tubes and somites in *Eed*-KO embryos, we refer to the primitive streak and NMPs as posterior tissues and uncover similar de-repression patterns (**Extended Data Fig. 4a**). Intriguingly, IM and LPM genes are not ectopically expressed in neural lineages, and brain genes do not upregulate in somitic tissues under PRC2 depletion (**Extended Data Fig. 4b-c**), which highlights that the ectopic de-repression is restricted to their corresponding tissue types.

To investigate if tissues respond differently to PRC2 depletion, we systematically analyzed differentially expressed genes in PRC2-depleted neural and somitic tissues through comparisons with their tissue-matched controls. Tissues with similar origins (i.e., neural tube 1 and 2 versus somite 0 and sclerotome) share common sets of upregulated genes as evidenced by the Jaccard similarity index^37^. (**Extended Data Fig. 4d**). By contrast, neural and somitic lineages share only a fraction of upregulated genes, which demonstrates that genes do not respond uniformly to PRC2 loss; instead, they manifest origin-specific sensitivity (**Fig. 2g**). GO analysis suggests that neural-specific upregulated genes are associated with brain development, whereas somitic-specific genes are associated with kidney and limb development (**Extended Data Fig. 4e-f**). To explore the tissue-specific responsiveness under PRC2 loss, we identified putative TFs that target neural and somite-specific upregulated genes, respectively^38^. We selected the top 25-ranked TFs from each gene set, filtered out non-expressing TFs in the control TLSs, and compared their expressions in neural tubes and somites using pseudobulk analysis (**Fig. 2h; Extended Data Fig. 5a-b**). TFs enriched only in the neural gene set show exclusive or high expression in neural tubes but are absent in somites, including *Sox1, Pou3f3, Sall3,* and *Gbx2*, which are important for neural development^39^. Similarly, TFs from the somite-specific gene set are uniquely or highly expressed in somites but not in neural tubes, including *Twist1/2, Foxc1/2, Prrx1,* and *Peg3,* which are essential for mesodermal development^40^. This indicates that gene-specific de-repression is conferred under the combinatory actions of (1) the loss of PRC2 activity, which facilitates a permissive transcription state, and (2) the presence of appropriate lineage-associated TFs to recruit the transcription machinery to targeted loci^41–43^. We conclude that the loss of PRC2 does not alter lineage commitment, but rather fosters an environment susceptible to stochastic gene de-repression, leading to increased transcriptional heterogeneity that compromises cellular fidelity. Additionally, we show that the full activation of ectopic lineage genes displays a TF-driven origin-specific sensitivity.

### Upregulated genes are defined by their chromatin states

To explore the role of chromatin more deeply, we profiled six epigenetic modifications in ESCs, control TLSs at 2, 3, 4, and 5 dpa using low-input Cut&Tag sequencing (**Fig. 1c**)^44^. We isolated neural (Venus-positive), mesodermal (mCherry-positive), and NMP (Venus/mCherry double-positive) cells from TLSs at 5 dpa using fluorescence-activated cell sorting (FACS), creating an *in vitro* embryoid chromatin landscape atlas with temporal and tissue-specific resolution. We included H3K27me3, H2AK119 ubiquitination (H2Aub), and H3K9 trimethylation (H3K9me3) as repressive chromatin marks; H3K27 acetylation (H3K27ac), H3K4me1, and H3K4me3 as associated with active/open chromatin marks^45^. We subsequently confirmed the expected enrichment patterns in ESCs, mesodermal, neural, and NMP lineages (**Extended Data Fig. 6a-b)**

To assess the lineage-specific H3K27me3 distribution, we performed peak calling and retrieved 6803 and 4038 H3K27me3 peaks in neural and mesodermal tissues, respectively (**Extended Data Fig. 6c**)^46^. The majority of mesoderm peaks overlap with neural lineage, indicating common genomic regions bound by PRC2. The enrichment of H3K27me3 also appears higher in neural cells at shared regions (**Extended Data Fig. 6d-g**). In contrast, the H2Aub levels are comparable in both lineages at the shared and neural-only H3K27me3 peaks, suggesting an autonomous recruitment of PRC1 as reported previously^47–49^.

Next, we systematically characterized chromatin features based on the combinatorial and spatial patterns of profiled histone marks using concatenated ChromHMM modeling. We defined a universal set of 15 common chromatin states shared across all samples while providing cell-type-specific annotations^50,51^. Among the 15 identified states are Polycomb repressed regions, bivalent transcription start sites (TSS), active TSS and active enhancers (**Fig. 3a**). We visually confirmed these state annotations with expected transition, including at the *Sox2* locus (**Fig. 3b**). We initially focused on chromatin state transitions of Cluster 5 and 8 genes to investigate whether gene clusters with differing temporal sensitivities exhibit distinct modules of upregulation. We used the mesodermal lineage as the end-stage reference due to its major representation. Most Cluster 5 genes stably remain as “bivalent TSS” or “repressed by Polycomb” throughout normal TLS development. This indicates that these genes should consistently remain silenced but are ectopically de-repressed under PRC2 loss (**Fig. 3c, Extended Data Fig. 7a**). Cluster 8 genes are initially active in the ESCs and a subset gradually becomes bivalent or repressed by Polycomb, while others progressively transition to quiescent states devoid of any profiled modifications, suggesting that these genes fail to silence following PRC2 loss (**Fig. 3d, Extended Data Fig. 7b**). The dynamic changes of H3K27me3, H2Aub, and H3K4me3 are consistent with the transitions in both clusters at locus-specific and global levels (**Fig. 3e-f, Extended Data Fig. 7c-d**). Our Cut&Tag data combined with ChromHMM annotations provides a comprehensive atlas of chromatin states and show how the chromatin environment evolves as cells transition from a pluripotent state toward fate specification and mechanistically underwrites the specific PRC2 dependence we observed.

### Transient PRC2 loss causes irreversible de-repression and phenotypic defects

Toward this end, we observed that PRC2 depletion during primitive streak induction generally led to more severe phenotypes. To take full advantage of the reversibility of the dTag system and explore the most sensitive windows of perturbation, we transiently depleted PRC2 for dT0-2 (during exit from pluripotency), dT2-3 (during PS induction), and dT0-3 by adding “Shield-1” after dTag^V-1^ washout to ensure rapid SUZ12 recovery (**Fig. 4a**)^20,52^. We first performed immunofluorescence (IF) and Cut&Tag to monitor the dynamics of SUZ12 and H3K27me3 after depletion or recovery. SUZ12 and H3K27me3 are reduced to levels comparable to those of *Suz12*-KO aggregates after 2 days of depletion (**Extended Data Fig. 8a-c**). Once Shield-1 is introduced, SUZ12 recovery can be observed as soon as 2 hours later and reaches a level similar to controls by 24 hours, while H3K27me3 show delayed and incomplete recovery (**Extended Data Fig. 8d-f**). We observe this delay in the other direction as well, with H3K27me3 showing a delayed reduction after the rapid PRC2 depletion (**Extended Data Fig. 8g-i**). Lastly, we also confirmed the reduction of H3K27me3 at its peaks under transient depletion (dT2-3, dT0-3; **Extended Data Fig. 8j-k**).

**Fig. 4:**
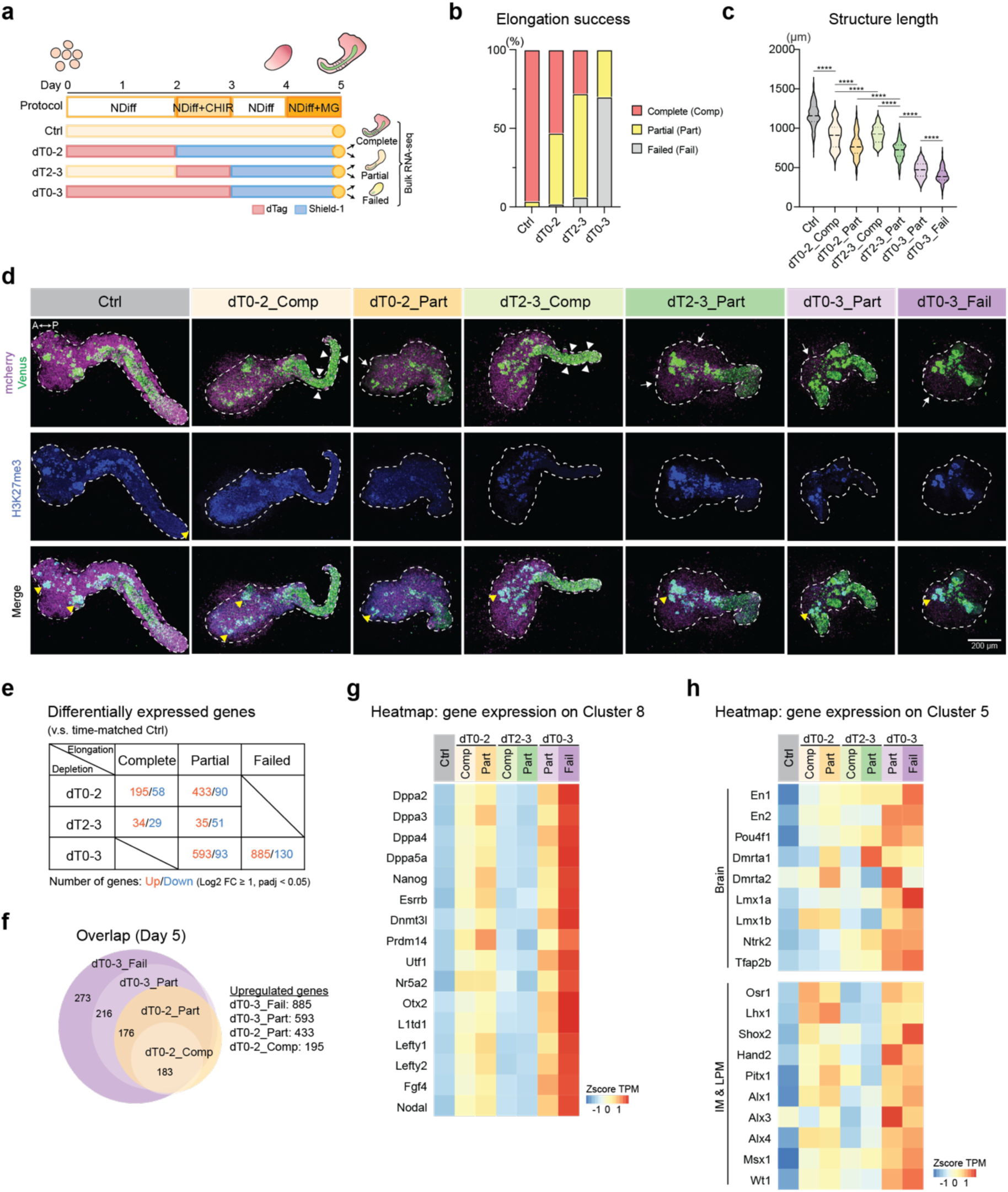
Transient PRC2 depletion causes irreversible de-repression and phenotypic defects. a, Schematic of transient PRC2 depletion conditions following sample collection for bulk RNA-seq. The red bars represent degradation periods, and the blue bars represent Shield-1 treatment, which leads to protein restoration. From each depletion condition, structures were sampled based on their phenotypes and perform bulk RNA-sequencing. **b,** Bar plot showing the percentage of elongation success. The structures were collected from two to four independent experiments (n ≥ 189 biological replicates). **c,** Violin plot displaying structure length across transient depletion conditions. The median is indicated by a dashed line, and the interquartile range is shown by dotted lines. This experiment was performed twice independently (n ≥ 44 biological replicates). **d,** Min-Max intensity projection of z-stacked TLSs confocal images. Anti-GFP, anti-mCherry and anti-H3K27me3 antibodies were multiplexed. White arrowheads indicate the observed lack of somitic tissues at the posterior end. White arrows display reduced or missing mCherry signal. Yellow arrowheads highlight the colocalization of GFP and H3K27me3 signals. The structures were outlined by dashed lines. n ≥ 5 biological replicates. **e**, Table showing numbers of differentially expressed genes across sample groups. **f,** Venn diagram representation of overlapping upregulated genes across sample groups. **g**, Heatmap displaying expression of Cluster 8 genes across sample groups. **h,** Heatmap analysis of Z-score normalized TPM of Cluster 5 genes across groups.

More than half of the TLSs subjected to the initial 2-day PRC2 depletion (dT0-2) are able to achieve complete elongation. However, by extending the depletion by just one day (dT0-3), most structures fail to elongate at all, and a fraction shows very limited protrusion. This striking effect underpins the requirement for PRC2 activity during WNT activation, which is further supported by the 24-hour depletion between day 2 and 3, causing pronounced morphological defects with more than 70% of the structures not elongating properly (**Fig. 4b, Extended Data Fig. 9a**). To further characterize the molecular dysregulation based on phenotypic severity, we categorized the TLSs into six sample groups by the depletion conditions (dT0-2, dT2-3, dT0-3) and by the morphological abnormalities (complete, partial, failed). Although reaching complete elongation, many TLSs still display abnormal phenotypes. They contain no or scarce somitic tissues at the posterior end, indicating that NMPs differentiate with a bias toward the neural lineage. In addition, mesodermal cells from partial or failed TLSs show smaller, fractured nuclei, indicating ongoing apoptosis, in accordance with observed tissue-specific lethality (**Fig 4c-d, Extended Data Fig. 9a**).

To interrogate the effects of perturbed transcriptional programs in PRC2 depleted conditions, we performed bulk RNA-seq and found that the scale of gene dysregulation positively correlates with the depletion duration as well as severity of phenotypic defects compared to control TLSs (**Fig. 4e-f, Extended Data Fig. 9b-c, Supplementary Table 2**). We previously linked differential temporal sensitivity to distinct modules of upregulation. Many Cluster 8 genes fail to silence upon early PRC2 depletion. As a subset gradually gain H3K27me3, we investigated whether restoring PRC2 activity could rescue this de-repression. Interestingly, if PRC2 is transiently absent during pluripotency exit (dT0-2 & dT0-3), many Cluster 8 genes are not able to be silenced even if PRC2 is restored afterwards (**Fig. 4g, Extended Data Fig. 9d**). However, if PRC2 depletion begins only after pluripotency exit, most Cluster 8 genes are not reactivated (dT2-3). These finding suggest they display a “window of sensitivity” where timely PRC2 activity is required. Once these genes are successfully silenced, they become inert to subsequent PRC2 depletion. In contrast, we observed that the brain, IM, and LPM genes, from the ectopically expressed Cluster 5 genes, are upregulated regardless of the depletion windows. This indicates that lineage-specific genes require continuous PRC2 presence; once de-repressed, the reestablishment of PRC2 activity does not rescue the misexpression. (**Fig. 4h, Extended Data Fig. 9e-f**). Lastly, genes associated with NMP identity and segmentation (Clusters 2 & 3) are consistently downregulated (**Extended Data Fig. 9f**).

Taken together, our dynamic PRC2 depletion conditions identified an increasing susceptibility to PRC2 perturbation during cell fate decisions, which can result in irreversible gene dysregulation and developmental defects.

## Discussion

Dynamic transcriptional activation and silencing ensure the establishment and maintenance of cellular identities. The lethality observed during gastrulation underscores PRC2’s critical role in regulating cell fate decisions^8^. Previous studies have shown that disrupting PRC2 prior to gastrulation already leads to abnormal epigenetic and cellular functions, emphasizing the necessity to target PRC2 loss directly during the gastrulation window^53–57^. Our TLSs system represents a reductionist approach to study early embryogenesis with high temporal resolution. It avoids the complexity of whole embryos, yet recapitulates the self-organization and lineage specification properties of developing embryos. Our findings revealed that PRC2 is required for pluripotency exit, consistent with a previous study showing retention of pluripotent factors under PRC2 loss during *in vitro* NPC differentiation^24^. However, through our temporally-resolved depletion scheme, we uncover that pluripotent genes only fail to silence with early depletion but are no longer reactivated with later depletion.

During TLS development, many genes exhibit irreversible transcriptional activation even under transient PRC2 perturbation. The unsuccessful reinstatement of repression potentially results from *trans*-acting, self-propagating TF networks which reinforce transcriptional activities, making silencing unattainable^58,59^. Interestingly, previous work in mESCs showed that reintroduction of PRC2 can accurately establish *de novo* H3K27me3 domains^60–62^, which highlights that this is possible in the steady-state pluripotency but as we show is no longer possible upon differentiation. This also fits recent work that showed a transient perturbation of PRC1 induces an irreversible cell fate switch in Drosophila^63^.

Genome-wide analysis of PRC1 and PRC2 occupancy and functional studies in mESCs suggest they act redundantly to repress genes^64,65^. However, our chromatin atlas in TLS reveals a unique enrichment of the PRC1 mediated H2Aub at many enhancers. Similarly, PRC1 binding at enhancers is observed in developing eye discs of Drosophila^66^. This implies an enhancer-dependent transcriptional control through PRC1 activity during embryogenesis, and a PRC1-mediated repression independent of PRC2, as reported in epidermal tissues^67^. As PRC1 inactivation (*Rnf2*-KO) is also embryonic lethal around gastrulation, it will be interesting to further explore how PRC1 and PRC2 safeguard embryogenesis^4,68,69^.

Together, we demonstrated an increasing and context-dependent reliance on the PRC2 repressive machinery during differentiation which improves our interpretation of the observed phenotypic defects. PRC2-mediated repression ensures that lineage-associated TFs act only upon intended targets, thereby preventing the stochastic activation of alternative developmental programs. Ultimately, our findings add to the understanding of core epigenetic mechanisms in governing cellular identities from pluripotency exit to lineage specification.

## Methods

### Mouse ESC culture

Mouse ESCs were cultured on 6-well plates coated with mitotically inactive primary mouse embryonic fibroblasts (MEFs) at 37°C and 5% CO₂. The culture medium consisted of KnockOut Dulbecco’s Modified Eagle Medium (DMEM, Gibco), supplemented with 15% fetal bovine serum (FBS, Gibco), 2 mM L-glutamax (Gibco), 100 U/ml penicillin-streptomycin (Gibco), 1× non-essential amino acids (Gibco), 1000 U/ml leukemia inhibitory factor (LIF), and 50 μM 2-mercaptoethanol (Gibco). Fresh medium was provided daily, and cells were resuspended with TrypLE (Gibco, 12604021) and split at a 1:10-15 ratio every 2 days.

### Generation of Suz12-dTag knock-in cells

The N-terminal of the Suz12 locus was targeted to incorporate the FKBP^F36V^ degradation tag by homology-directed repair (HDR) using the Alt-R CRISPR-Cas9 genome editing system (IDT)^20,21^. Briefly, CRISPR RNA (crRNA) targeting *Suz12* was designed using the CRISPOR tool and synthesized by IDT^70^. The crRNA was then annealed with tracrRNA and incubated with Cas9 nuclease (IDT, 1081059) to assemble the CRISPR-Cas9 ribonucleoprotein (RNP) complex. Homology arms flanking the targeted sequence were amplified by Q5 high-fidelity DNA polymerase (NEB) and cloned into the pCRIS-PITChv2-Puro-dTAG plasmid (Addgene #91793). The repair template containing the homologies, puromycin resistance, P2A peptides, and dTag cassettes was then amplified by PCR and co-delivered with the RNP complex into mESCs using electroporation with the Nucleofector System (Lonza, V4XP-3024). Cells were then recovered for 48 hours in culture media supplemented with Alt-R™ HDR Enhancer (IDT, 10007910). After 5 days of puromycin selection (Thermo, A1113803), single-cell-derived homozygous clones were confirmed by genotyping using the Phire Tissue Direct PCR Master Mix (Thermo, F170L).

### Generation of trunk-like structures

Cells were passaged at least two times after thawing, and TLSs were generated as previously described with minor modifications^16^. After cell suspension, the mESC population was enriched using feeder removal microbeads (Miltenyi, 130-095-531). After centrifugation, mESCs were resuspended in NDiff medium (Takara, XA0530), and 500 cells (in 40 μl of cell suspension) were seeded in a 96-well ultra-low attachment round-bottom plate (Corning, 7007). At 2 days post-aggregation, 150 μl of NDiff medium supplemented with 3 μM CHIR99021 (Sigma, SML1046) was added to the well. At 3 dpa, 150 μl of fresh medium was replaced. At 4 dpa, the medium was changed, and 150 μl of NDiff medium with 5% Matrigel (Fisher, 356231) was added. At 5 dpa, the TLSs were subjected to downstream analysis. The compounds dTagV-1 (Tocris, 6914) or dNeg (Tocris, 6915) were diluted to reach a final concentration of 500 nM.

### PRC2 restoration by washout experiments

For washout experiments, aggregates were briefly washed by sequentially removing and adding 150 μl of pre-warmed media twice. Afterward, 150 μl of media with 500 nM Shield-1 (Takara, 632189) and the appropriate components (i.e., CHIR or Matrigel) was added.

### Morphometric analysis of TLS

For scoring of axis elongation, TLSs at 5 dpa were imaged using Zeiss CellDiscoverer 7. Structures were categorized based on elongation status including “Complete”, “Partial”, “Failed”, and “Multiaxes”. If a structure displayed no protrusion, it was scored as “Failed” elongation. If a structure showed minor to mild protrusion, it was scored as “Partial” elongation. If a structure reached complete elongation but displayed two or more axes, it was scored as “Multiple axes”. To quantify structure length, the longest path was measured. For structures with multiple axes, the longest path containing a single axis was used. For statistical analysis, sample groups were assessed for normality using the Shapiro-Wilk test. Statistical significance was determined via one-way ANOVA followed by Tukey’s honest significant difference (HSD) test for multiple comparisons. For stable depletion experiments, *F* = 199.0 with degrees of freedom (DF) = 5. For transient depletion experiments, F = 510.4 with DF = 6. All conditions represent at least two independent experiments. For all conditions, at least two independent experiments were performed.

### Western blot

Samples were lysed with RIPA buffer (Thermo, 89900) supplemented with protease inhibitors (Thermo, 78443). Protein concentration was determined by the Bradford assay. Five to thirty μg of protein extract was used for electrophoresis using a pre-cast gradient gel (Thermo). The samples were transferred to nitrocellulose membranes (Bio-Rad). Membranes were blocked for 1 hour with 5% skimmed milk in TBST, then incubated with primary antibodies diluted at a 1:1000 ratio in TBST overnight at 4°C with gentle shaking. After three washes with TBST, membranes were incubated with fluorophore-conjugated secondary antibodies (1:10000) at room temperature in the dark. After another three washes, the images were developed using the ChemiDoc Imaging System (Bio-Rad).

### Bulk RNA-sequencing

Cells or aggregates were collected, washed twice with PBS, and snap-frozen in liquid nitrogen. RNA was extracted using the RNeasy Plus Micro Kit (Qiagen, 74034) following the manufacturer’s instructions. RNA concentration was measured using the Qubit RNA High Sensitivity Assay (Invitrogen, Q32852). Samples with a RIN number greater than 8.0 (Agilent TapeStation) proceeded to library preparation (KAPA) and sequencing with NovaSeq.

### Single cell RNA-sequencing

Single-cell RNA sequencing experiments were performed using the Parse Biosciences Evercode WT kit following the manufacturer’s instructions. Briefly, a single-cell suspension was acquired by incubation with TrypLE at 37°C for 30 minutes, washed twice with PBS, and filtered through a 40 μm cell strainer (Falcon) to remove cell debris and clumps. After gentle fixing, fixed cells were snap-frozen in liquid nitrogen and stored at-80°C. Plate-based barcoding and library amplification were performed following the manufacturer’s instructions. The quality of the library was determined by Agilent TapeStation analysis, and Illumina-based sequencing was performed by Parse Biosciences with 200 million reads.

### Cut&Tag-sequencing

Low-input Cut&Tag was performed as previously described with minor modifications^44^. TLSs were pooled to minimize structure-to-structure variations. Briefly, aggregates were dissociated by incubation with TrypLE at 37°C for 30 minutes. To acquire lineage-specific resolution in Day 5 TLSs, the cell suspension was subjected to FACS to isolate Venus-positive, mCherry-positive, and double-positive populations representing neural, mesodermal, and NMP cells, respectively. Ten thousand cells were used per epitope, and the total number of cells was scaled up accordingly. Cells were suspended in wash buffer and incubated with Concanavalin A beads (Biomag) at room temperature for 10 minutes with gentle rotation (10 μl beads per reaction, scaled up accordingly). Cells were resuspended in 50 μl of antibody buffer, and 0.5 μl of primary antibody was added. After overnight incubation at 4°C with gentle rocking, cells were washed twice with Dig-wash buffer and incubated with secondary antibody (1:100 dilution in antibody buffer) for 60 minutes at room temperature with gentle rocking. After two washes with Dig-wash buffer, cells were incubated with pA-Tn5 (1:400 dilution, Institute Curie) for 1 hour at room temperature. After three Dig-wash buffer washes, cells were resuspended with 100 μl of tagmentation buffer supplemented with yeast spike-in DNA (20 ng/μl, Cell Signaling Technology) and incubated at 37°C for 1 hour. Afterward, EDTA, SDS, and proteinase K were added and incubated at 55°C for 1 hour. DNA was purified using a DNA clean-up kit (Zymo). For library preparation, 21 μl of DNA was mixed with 2 μl of uniquely barcoded i5 primer, 2 μl of uniquely barcoded i7 primer, and 25 μl of 2× PCR Master Mix (NEB). The PCR program was set as follows: 72°C for 5 minutes; 98°C for 30 seconds; 14 cycles of 98°C for 10 seconds and 63°C for 30 seconds; final extension at 72°C for 1 minute, and samples were held at 4°C. Post-PCR clean-up was performed by Ampure XP beads (Beckman Coulter) and size distribution was determined by Agilent TapeStation analysis. Sample libraries were pooled to reach equal representation and subjected to paired-end sequencing following the manufacturer’s instructions.

### Whole mount Immunofluorescence

For TLSs developed ranging from 2-3 dpa, samples were collected into 2 ml protein low-binding tubes (Eppendorf, 0030108132) pre-coated with PBS containing 10% FBS. Centrifugation was performed in a swinging-bucket rotor at 10 g for 2 minutes at room temperature. After two PBS washes, samples were fixed with freshly prepared 4% PFA at room temperature for 30 minutes. After three PBS washes, samples were permeabilized with 0.5% Tween-20 in PBS (PBST) at room temperature for 30 minutes and incubated overnight with blocking buffer [3% donkey serum (Biozol, LIN-END9010) in PBST] at 4°C with gentle shaking. Subsequently, aggregates were incubated overnight with primary antibodies diluted 1:200 in blocking buffer at 4°C with gentle shaking. After three washes with blocking buffer, samples were incubated overnight at 4°C with secondary antibody (1:250 dilution, Jackson Labs) and DAPI (0.2 μg/ml, Thermo Scientific, 62248). After two PBS washes, aggregates were resuspended with Vectashield mounting medium (VectorLabs, H-1900), transferred to glass chamber slides, sealed with coverslips, and stored in the dark at 4°C.

The procedure for TLSs developed ranging from 4-5 dpa was similar, except the samples were transferred to μ-well glass-bottom coverslips (ibidi, 80827-90). After fixation, permeabilization, and overnight blocking, aggregates were incubated with primary antibody at 4°C for 48-72 hours. After three washes with blocking buffer, TLSs were incubated overnight at 4°C with secondary antibody (1:250 dilution, Jackson Labs) and DAPI (0.2 μg/ml, Thermo Scientific, 62248). After two washes with PBS and two washes with 0.02 M phosphate buffer (PB), TLSs were incubated in RIMS buffer [133% Histodenz (Merck, D2158) in 0.02 M PB] at 4°C overnight for tissue clearing.

### Microscopy

TLSs stained with antibodies were imaged using a Zeiss LSM880 confocal laser scanning microscope with Airyscan, using filters for Alexa Fluor 488, Alexa Fluor 594, Alexa Fluor 647, DAPI, and combinations thereof. Images were processed and analyzed using Zen and Fiji. For live imaging, TLSs were imaged with a Zeiss CellDiscoverer 7, with the incubator chamber temperature set at 37°C and CO₂ content at 5%. Acquisition was set for brightfield, mCherry, and Venus with z-stacking set to 10 μm.

### Bulk RNA-seq data preprocessing and data analysis

Raw reads were subjected to adapter and quality trimming with cutadapt (version 4.4; parameters: --quality-cutoff 20 --overlap 5 --minimum-length 25 --interleaved --adapter AGATCGGAAGAGC-A AGATCGGAAGAGC), followed by poly-A trimming with cutadapt (parameters: --interleaved --overlap 20 --minimum-length --adapter “A[100]” --adapter “T[100]”). Reads were aligned to the mouse reference genome (mm10) using STAR (version 2.7.9a; parameters: --runMode alignReads --chimSegmentMin 20 --outSAMstrandField intronMotif --quantMode GeneCounts) and transcripts were quantified using stringtie (version 2.0.6; parameters:-e) with GENCODE annotation (release VM19). Differential gene expression testing has been done with DESeq2 (version 1.44.0). Heatmaps were created with the ComplexHeatmap package in R.

### CUT&Tag data preprocessing and analysis

CUT&Tag data processing and analysis were done with the cutandrun nf-core pipeline (doi:10.5281/zenodo.5653535). In particular, reads were aligned to mouse reference genome GRCm38. Yeast (*S. cerevisiae*) DNA was used for spike-in normalization. Peaks were called with MASC2^46^, and consensus peaks between two replicates were extracted for downstream analysis. Metaplots and heatmaps were generated using Deeptool^71^. Chromatin states were determined using ChromHMM^50,51^. In particular, bedGraph output files from the cutandtag pipeline were first binned so that every condition has the same bin coordinates. Next, files were combined per chromosome and then ChromHMM was run with the BinarizeSignal command.

Last, a model with 15 chromatin states was created by running ChromHMM with the LearnModel command. Figures have been created with custom R scripts from the model output files. Alluvial plots were created with ggplot2 using the state emission probabilities of the TSS.

### Single-cell RNA-seq data preprocessing and analysis

FASTQ files were processed using the Trailmaker pipeline module (https://app.trailmaker.parsebiosciences.com/; pipeline v1.5.1, Parse Biosciences). Unfiltered count matrices were imported into R (v4.4.1) and analyzed with Seurat (v5.3.0)^33^. Cells with fewer than 1000 detected genes, more than 100,000 counts (controls) or 75,000 counts (degron samples), or assigned to undefined clusters were excluded. Cell cycle phases were annotated with mouse orthologs of the S and G2M gene sets using Seurat’s CellCycleScoring.

Samples were integrated with Harmony (theta = 2) based on PCA embeddings of the 2000 most variable genes^34^. UMAPs were computed from the first 16 Harmony dimensions, and clustering was performed at a resolution of 2.4 using Louvain clustering. Cell types were assigned to clusters by manual inspection of marker gene expression.

We quantified the cellular heterogeneity by computing the average pairwise Euclidean distance among cells within each sample and cell type in the Harmony space. Harmony-aligned embeddings for the first 16 dimensions were extracted and filtered for Neural tube 1, Neural tube 2, and Somite (Sclerotome) cells from all t= day 5 timepoint samples. For each group, we performed a bootstrap procedure. Only groups containing at least 20 cells were included. For every eligible group, we repeatedly drew subsamples of 35 cells, and sampling was performed without replacement. For each bootstrap iteration, we calculated the Euclidean distance matrix of the subsampled cells and recorded the mean of all pairwise distances. This procedure was repeated five hundred times per group, yielding a distribution of average pairwise distances. Group differences were tested by one-way ANOVA with Tukey’s post hoc test (stats package). Visualizations used ggplot2 (v3.5.2).

Differential gene expression was assessed using Seurat’s FindMarkers, comparing depleted samples to the Day 5 control (min.pct = 0.1, adjusted p < 0.05). Cell type similarities were evaluated using the Jaccard index on sets of differentially expressed genes and visualized with ComplexHeatmap (v2.20.0)^37^. Transcription factor enrichment in genes belonging to the neural-specific, somit-specific, or shared gene lists was determined using ChEA3. Pseudobulk samples were generated using Seurat’s AggregateExpression function, grouping Neural tube 1 and Neural tube 2 cells together as Neural and Somite (0) and Somite (Sclerotome) cells as Somite. TPMs were calculated by extracting the exon lengths from RefSeq to calculate the transcript length for the longest transcript, and the convertCounts function from DGEobj.utils (v1.0.6). Log fold-changes were determined for any of the top 25 ChEA3 transcription factors that showed strong expression in the scRNA data^38^. Expression at the single-cell level was determined using the VlnPlot function from Seurat.

## Acknowledgements

We thank all members of the A. Meissner laboratory for their critical input. We also thank M. Awawdy for help with image analysis. Our gratitude extends to A. Balaskas and A. Bulut-Karslioğlu for scientific discussions. We are grateful to the Sequencing Core Facility and M. Piedavent-Salomon of the Flow Cytometry Facility for their support. M.-K. Lee is a fellow of the European Union’s HORIZON-MSCA-2022 postdoctoral fellowship (Project ID 101109517, DyMERE). Funding was provided by the European Union (ERC, CancerEpigenome, 101098178). Views and opinions expressed are however those of the author(s) only and do not necessarily reflect those of the European Union or the European Research Council. Neither the European Union nor the granting authority can be held responsible for them. This work is generally supported by the Max Planck Society.

## Contributions

M.-K.L. and A.M. designed and conceived the study. M.-K.L., S.D.M., and A.M. prepared the manuscript with the assistance of the other authors. M.-K.L. generated the SUZ12-dTag cell line, performed the TLSs experiments, and all the sample preparation and libraries preparation for bulk RNA-seq, scRNA-seq, and CUT&Tag-seq. M.-K.L. performed the staining, imaging, and image analysis. S.D.M. performed the bulk RNA-seq, scRNA-seq and Cut&Tag analyses.

J.V. performed the scRNA-seq and Cut&Tag analyses. D.F. performed the Western blot analysis and live cell imaging. M.W. helped with the bulk RNA extraction. A.M. supervised the work.

## Competing interests

The authors declare no competing interests.

## Extended Data

**Extended Data Fig. 1:**
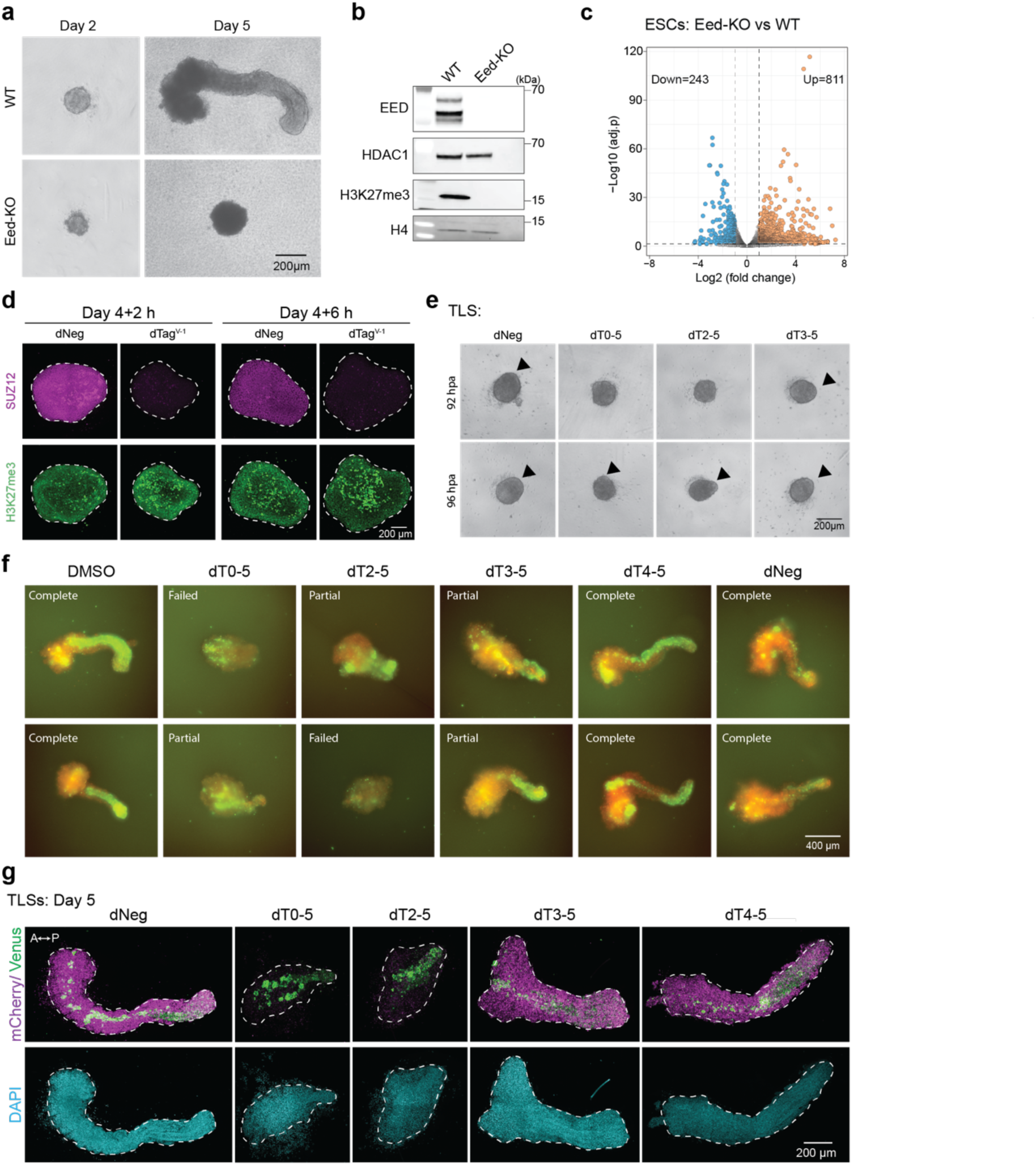
PRC2 inactivation causes failed TLS development. **a,** Bright field images of WT and *Eed*-KO TLSs at 2 and 5 dpa. **b,** Western blot analysis showing protein levels of EED and H3K27me3 in WT and *Eed*-KO mESCs. HDAC1 and H4 were used as loading controls. **c,** Volcano plot displaying differentially expressed genes in *Eed*-KO cells compared to WT mESCs. Each dot represents a single gene. Orange dots indicate significantly upregulated genes, and blue dots indicate significantly downregulated genes, based on a threshold of Log2 fold change (Log2FC) ≥ 1 and *P*_adj_ < 0.05. **d,** Confocal microscopy showing the abundance of SUZ12 and H3K27me3 under dTag treatment. **e,** Bright field images of TLSs at 92 and 96 hpa under PRC2 depletion. The arrowheads indicate the protrusion of an elongating front. **f,** Live-cell imaging showing representation of complete, partial, failed elongation. **g,** Whole-mount IF of TLSs. Anti-GFP and anti-mCherry antibodies were used to stain neural and mesodermal tissues, respectively. Dashed lines represent the contour of the TLSs.

**Extended Data Fig. 2:**
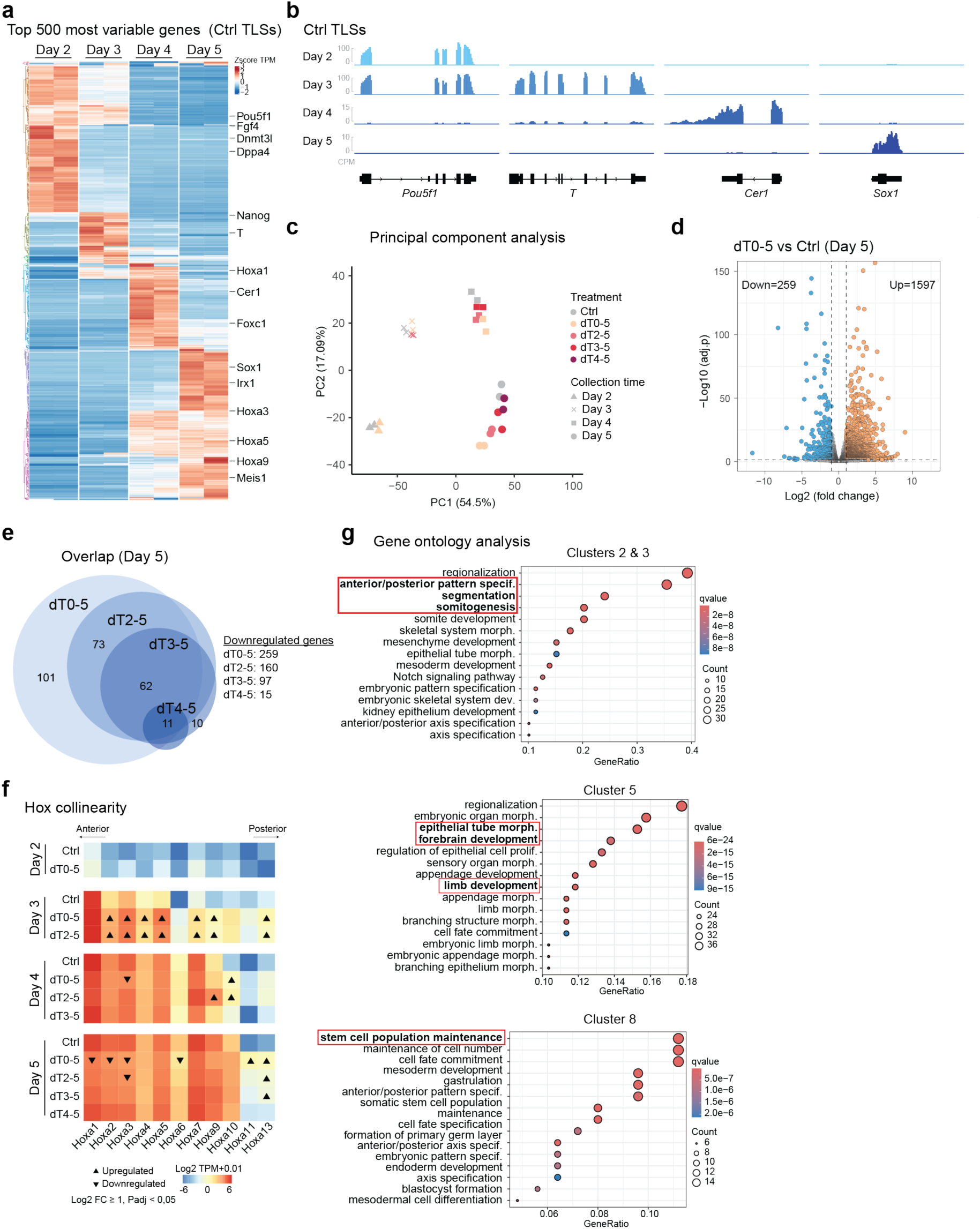
Transcriptomic analysis of developing TLSs under PRC2 loss. **a,** Heatmap analysis showing the top 500 most variable genes across control TLSs at 2, 3, 4 and 5 dpa. Clustering was performed using Euclidean distance metrics. Color intensity represents Z-score normalized TPM levels. **b,** Genomic snapshots displaying RNA-seq in control TLSs across time points. CPM, counts per million. **c,** Principal component analysis of TLSs across depletion conditions and time points. The colors denote depletion conditions. The shapes represent sample collection time points. **d,** Volcano plot displaying differentially expressed genes dT0-5 versus Ctrl at 5 dpa. **e,** Venn diagram showing the overlap of downregulated genes across depletion conditions in TLS at 5 dpa. **f,** Heatmap analysis of *Hoxa* locus expression represented in Log2 TPM+0.01. The value represents the mean of two biological replicates. **g,** Gene ontology analysis of genes from Clusters 2 and 3, Cluster 5, and Cluster 8. The top 15 ontologies were selected.

**Extended Data Fig. 3:**
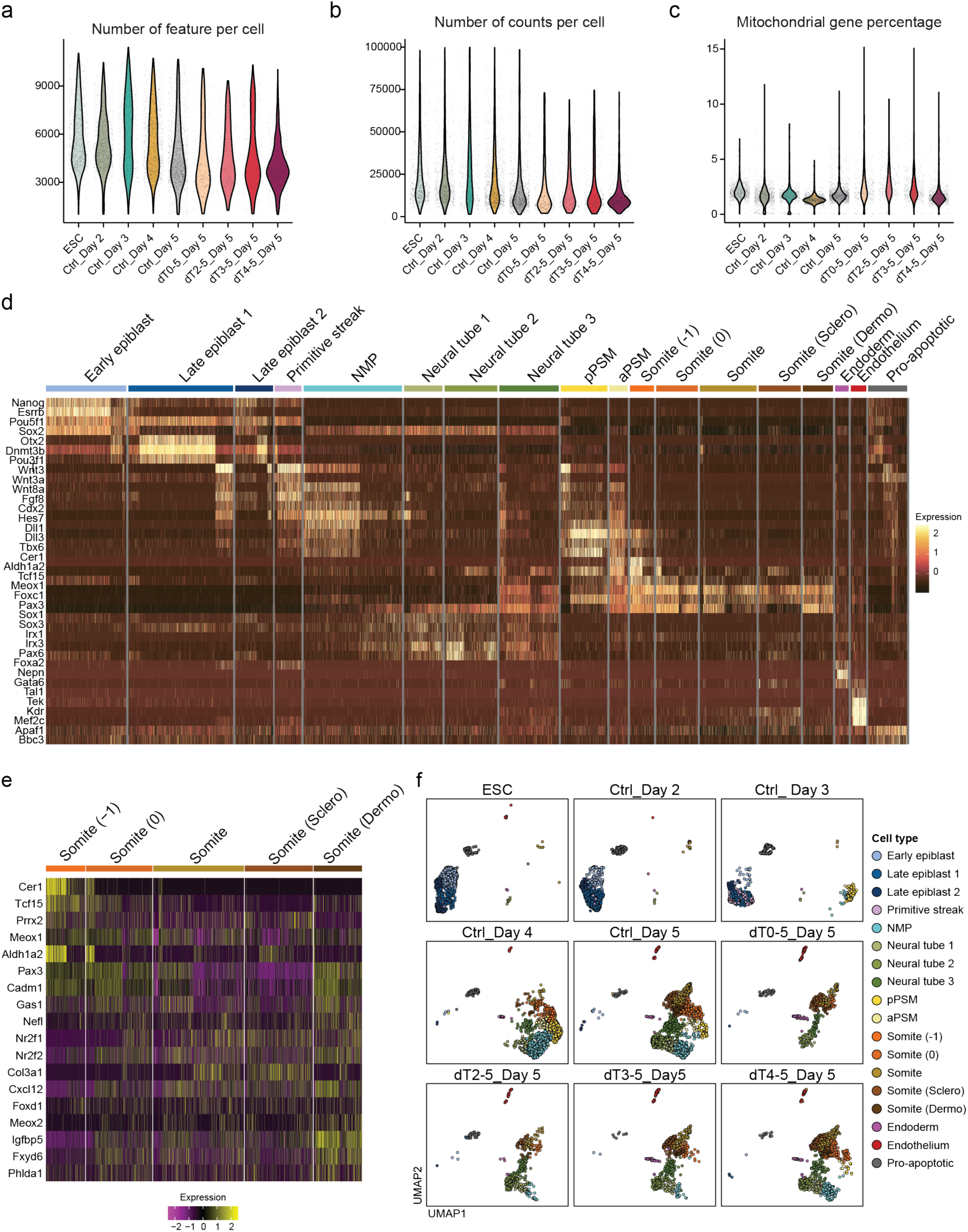
Single cell-RNA analysis of PRC2-depleted TLSs. **a,** Violin plot showing the number of features per cell across samples. **b,** Violin plot displaying number of counts per cell across samples. **c,** Violin plot showing the percentage of mitochondrial genes number per cell across samples. **d,** Heatmap analysis showing scaled expression of genes associated with pluripotency, primitive streak, NMP, somitogenesis, and neural tubes measured in 9091 cells across cell types. **e,** Heatmap analysis showing scaled expression levels of genes associated with somitogenesis. **f,** Dimensional reduction plots showing representation of cell types across time points and depletion conditions.

**Extended Data Fig. 4:**
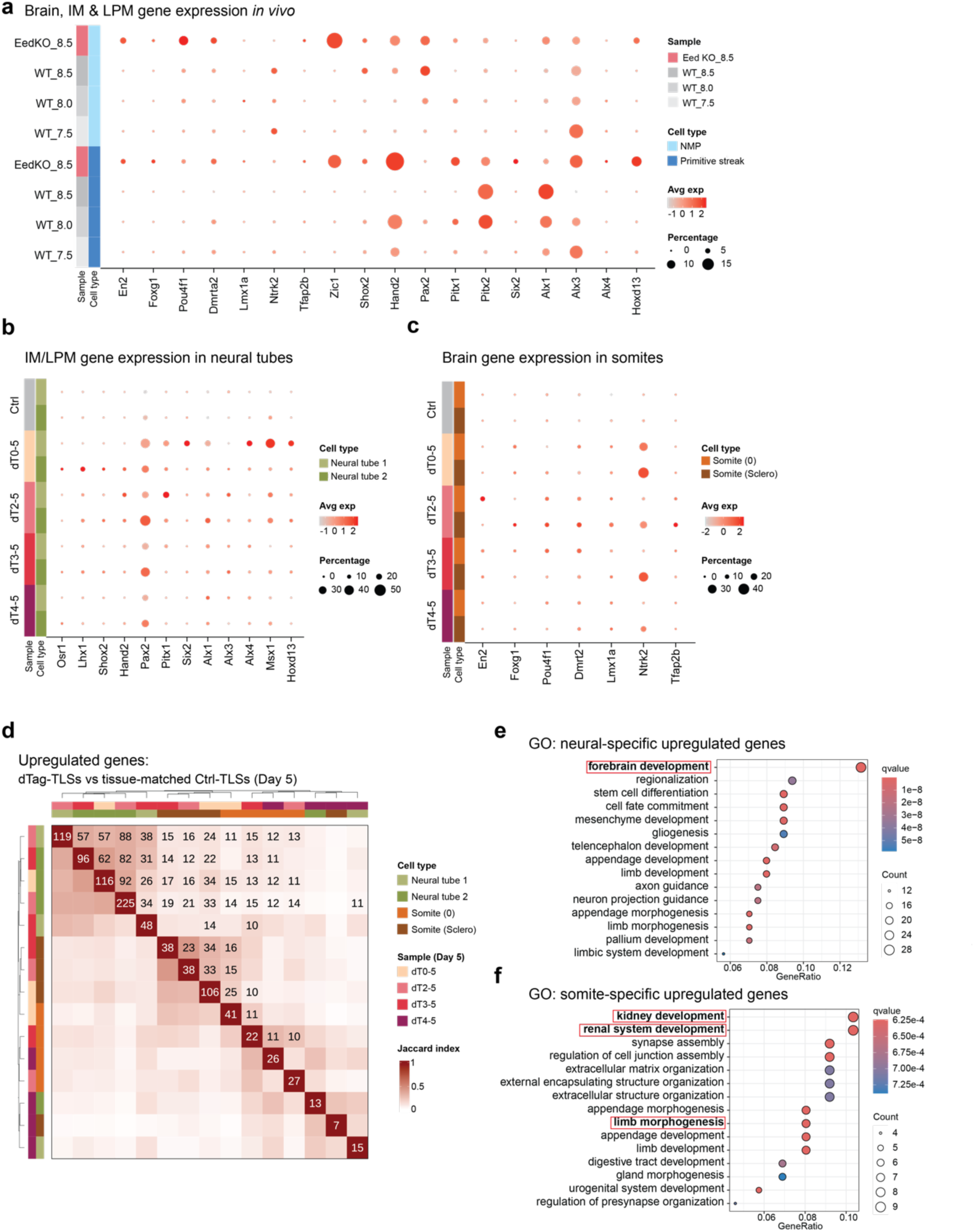
Single-cell RNA analysis of deregulated genes across depletion conditions. **a,** Dot plot showing the expression levels of brain, IM, and LPM markers in *Eed*-KO (E8.5) and WT mouse embryos (E7.5-8.5). Primitive streak and NMPs were represented as posterior tissues. **b,** Dot plot displaying expression of IM/LPM markers in neural tube 1 and 2 across samples at 5 dpa. **c,** Dot plot representation of the expression of brain markers in somite (0) and somite (sclerotome) across depletion conditions at 5 dpa. **d,** Heatmap showing the overlap of upregulated genes across cell types and depletion conditions. Differential analysis was performed with tissue-matched control cell type. Neural tube 1 and 2, somite (0), and somite (sclerotome) were compared. The tissue neural tube 1 from group dT0-5 was excluded from the analysis due to scare cell number (n = 5). The similarity is analyzed by Jaccard index, representing by intensity of the color. The values in white denote the number of upregulated genes in the sample. The values in black represent the number of overlapped genes between samples. For visualization and clarity, overlapped numbers below 10 were not shown. **e,** Gene ontology analysis of neural-specific upregulated genes as in Fig. 2g. **f,** Gene ontology analysis of somite-specific upregulated genes as in Fig. 2g.

**Extended Data Fig. 5:**
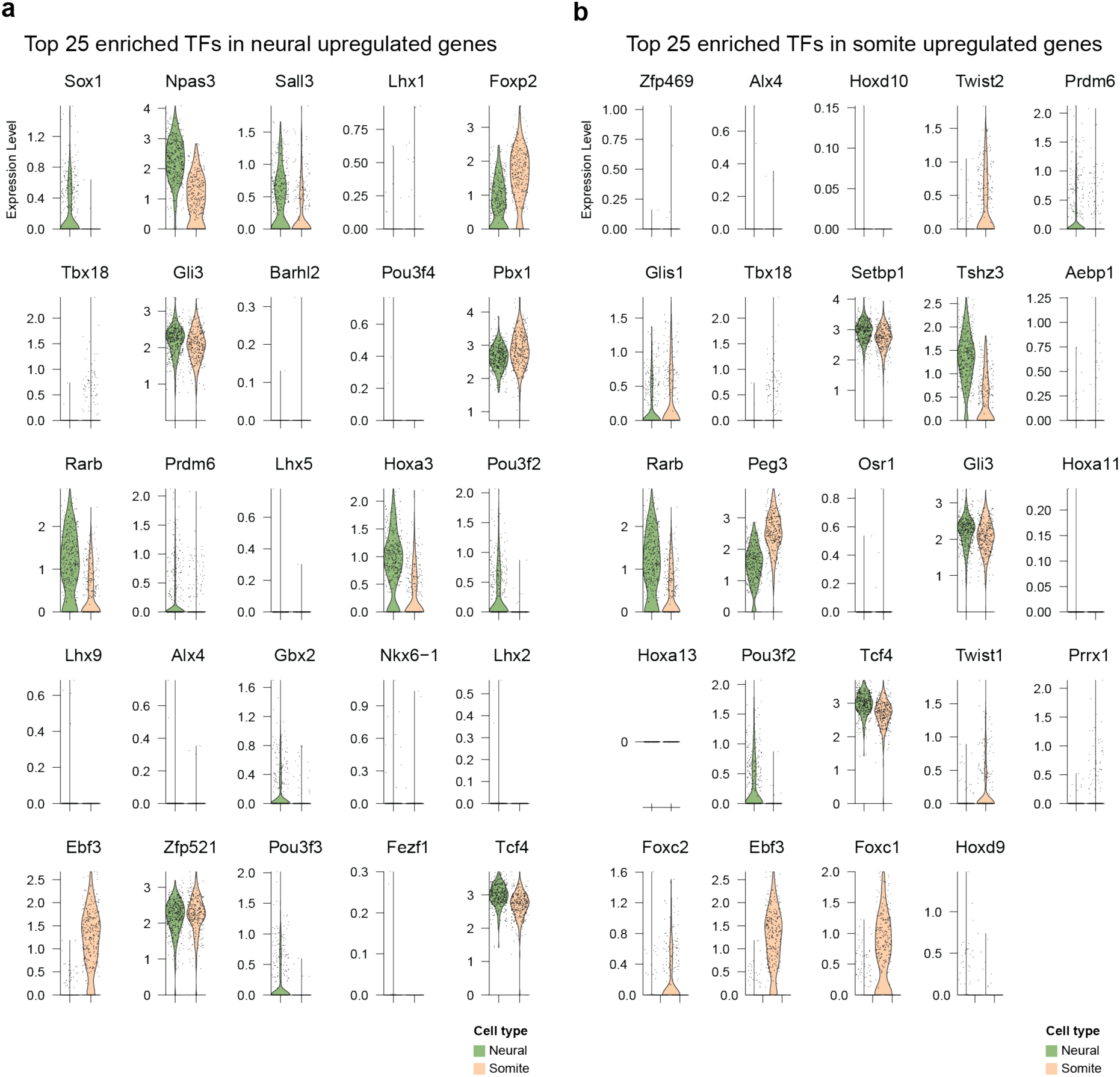
Single-cell transcriptomic analysis of enriched TFs expression. **a,** Violin plot showing the expression of the top 25 enriched TFs of neural-specific upregulated genes in neural lineages (neural tube 1 and 2) and mesodermal lineages (somite 0 and sclerotome). **b,** Violin plot showing the expression of the top 25 enriched TFs of somite-specific upregulated genes in neural lineages (neural tube 1 and 2) and mesodermal lineages (somite 0 and sclerotome).

**Extended Data Fig. 6:**
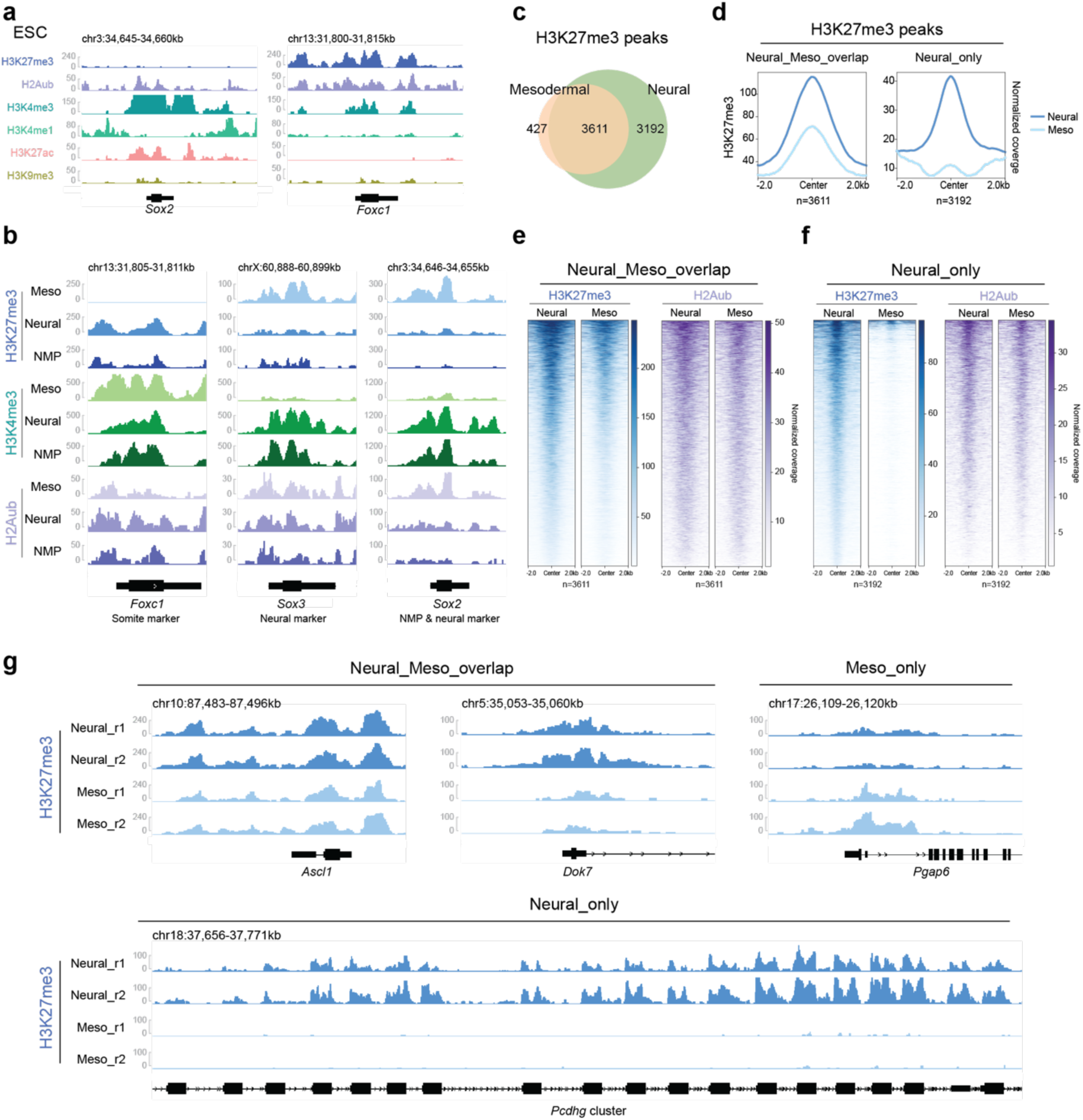
Cut&Tag-sequencing captures chromatin landscapes.**a,** Genomic snapshots showing the levels of H3K27me3, H2Aub, H3K4me3, H3K4me1, H3K27ac, and H3K9me3 at the *Sox2* and *Foxc1* loci in mESCs. **b,** Genomic snapshots displaying the levels of H3K27me3, H2Aub, H3K4me3 in mesodermal cells, neural cells and NMPs at the *Foxc1, Sox3* and *Sox2* loci. **c,** Venn diagram representation of overlapping H3K27me3 peaks in neural and mesodermal lineages. **d,** Metaplot analysis of H3K27me3 and H2Aub levels at neural-specific peaks and overlapping peaks in neural and mesodermal lineages as represented in **c**. **e,** Heatmap analysis of H3K27me3 and H2Aub enrichment at overlapping H3K27me3 peaks in neural and mesodermal tissues. **f,** Heatmap showing levels of H3K27me3 and H2Aub at neural-specific H3K27me3 peaks in neural and mesodermal tissues. **g,** Genomic snapshots displaying examples of H3K27me3 distribution at overlapping, mesodermal-specific, and neural-specific H3K27me3 peaks in neural and mesodermal tissues.

**Extended Data Fig. 7:**
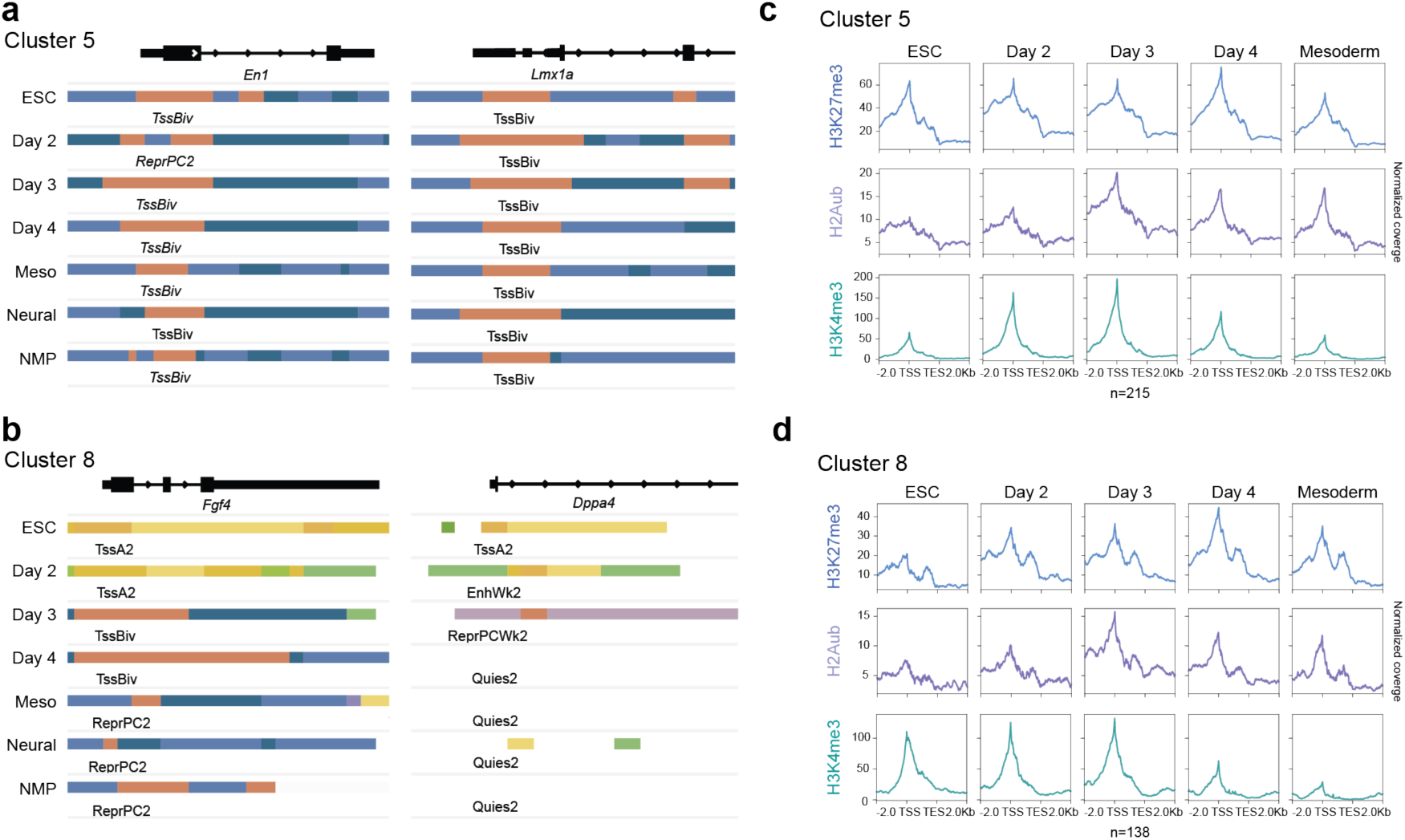
Chromatin state transitions revealed by ChromHMM. **a,** Genomic snapshot showing the chromatin state transition of the *En1* and *Lmx1a* locus, as examples of Cluster 5 genes. **b,** Genomic snapshot showing the chromatin state transition of the *Fgf4* and *Dppa4* locus, as examples of Cluster 8 genes. **c,** Metaplot analysis displaying levels of H3K27me3, H2Aub and H3K4me3 in Cluster 5 genes across samples. **d,** Metaplots showing H3K27me3, H2Aub and H3K4me3 levels in Cluster 8 genes across samples.

**Extended Data Fig. 8:**
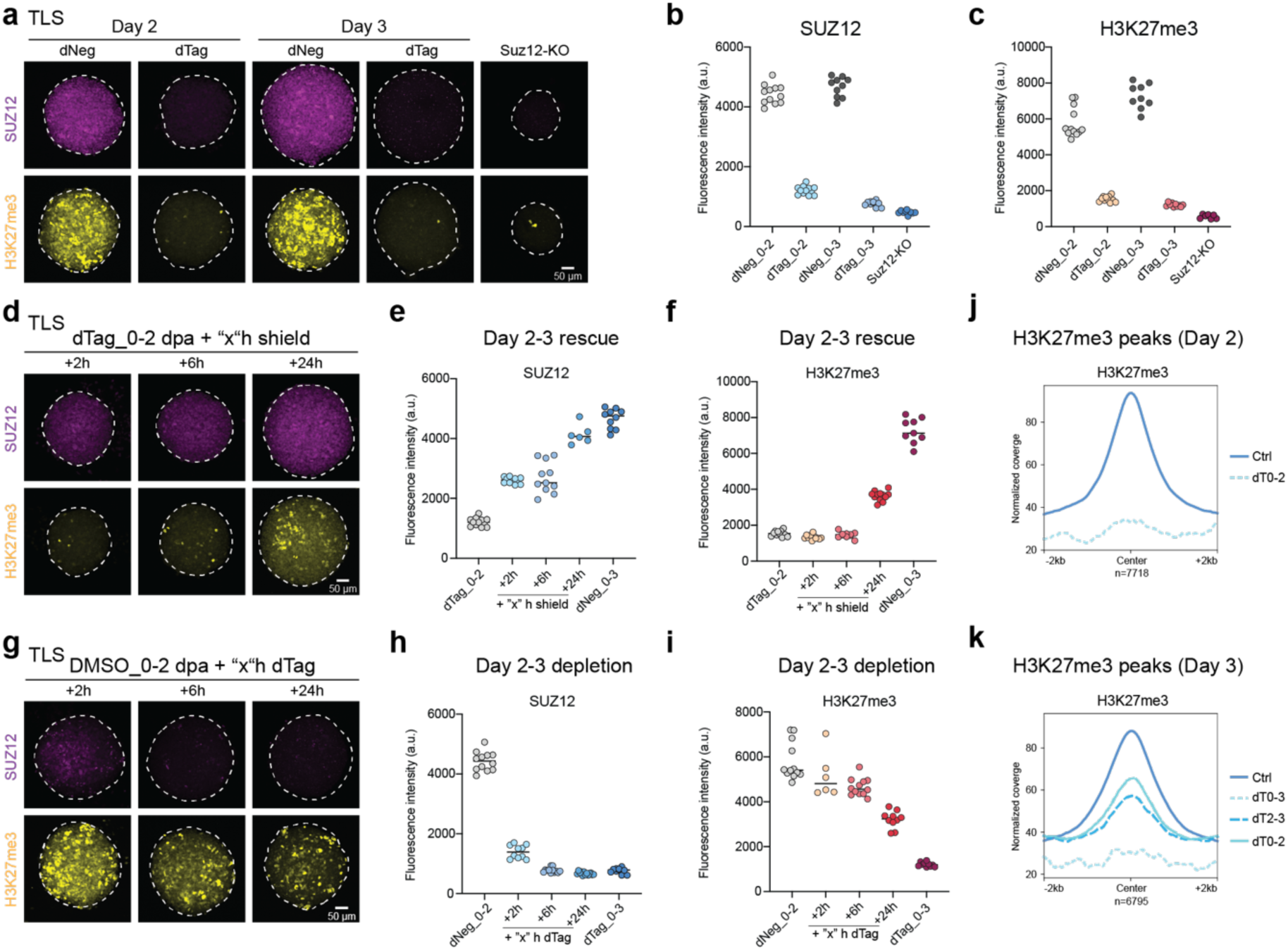
Dynamics of PRC2 and H3K27me3 upon degradation and restoration. **a,** Maximum intensity projection of z-stacked IF staining of SUZ12 and H3K27me3 in TLSs at 2 and 3 dpa, and Suz12-KO derived TLSs. **b,** Arbitrary fluorescence intensity of SUZ12 level from data shown in **a**. n ≥ 9 biological replicates. **c,** Arbitrary fluorescence intensity of H3K27me3 from data shown in **a**. n ≥ 9 biological replicates. **d,** Maximum intensity projection of z-stacked IF staining of SUZ12 and H3K27me3 of PRC2-depleted TLSs followed by Shield-1 treatment to allow PRC2 restoration. TLSs were treated with dTag for 2 days, and samples were collected at 2, 6 and 24 hours after Shield-1 treatment. **e,** Arbitrary fluorescence intensity of SUZ12 from data shown in **d**. n ≥ 6 biological replicates. **f,** Arbitrary fluorescence intensity of H3K27me3 from data shown in **d**. n ≥ 6 biological replicates. **g,** Maximum intensity projection of z-stacked IF staining of SUZ12 and H3K27me3 in TLSs followed by PRC2 degradation. After 2 days post-aggregation, TLSs were collected at 2, 6 and 24 hours after dTag treatment. **h,** Arbitrary fluorescence intensity of SUZ12 from data shown in **g**. n ≥ 10 biological replicates. **i,** Arbitrary fluorescence intensity of H3K27me3 from data shown in **g**. n ≥ 10 biological replicates. **j,** Metaplot analysis of H3K27me3 levels in control TLSs and TLSs under 2 days of dTag treatment (dT0-2). The signals were plotted over H3K27me3 peaks in ctrl TLSs at 2 dpa. **k,** Metaplot analysis of H3K27me3 levels across depletion conditions TLS at 3 dpa. The signals were plotted over H3K27me3 peaks in ctrl TLSs at 3 dpa. The samples included control TLSs, TLSs under 3 days of depletion (dT0-3), TLSs under 2 days of depletion following 1 day of Shield-1 recovery (dT0-2), and TLSs under 1 day of depletion (dT2-3).

**Extended Data Fig. 9:**
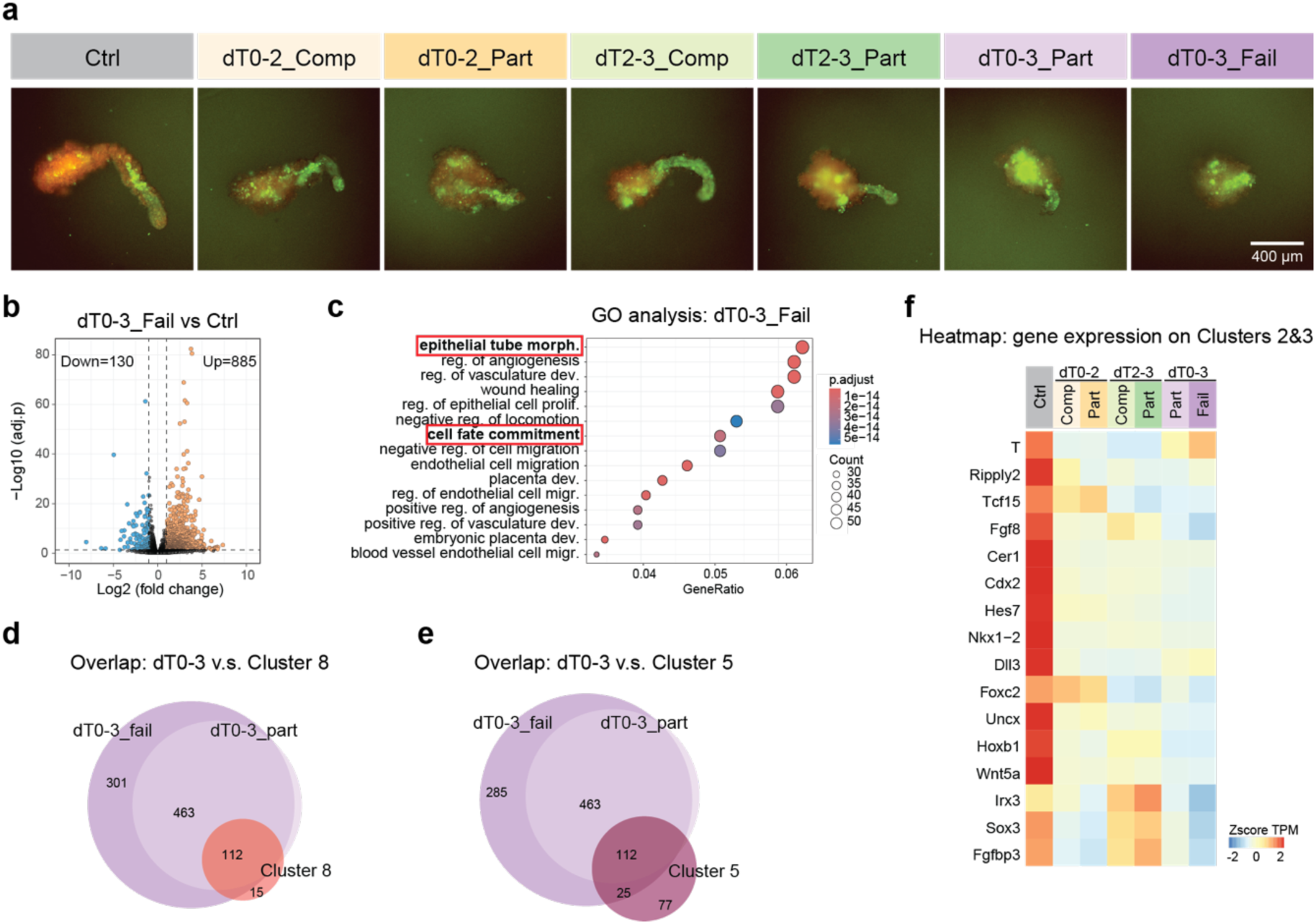
Transient PRC2 depletion perturbs differentiation programs. **a,** Live-cell images of TLSs across sample groups. **b,** Volcano plot displaying differentially expressed genes in dT0-3_failed versus ctrl. **c,** Gene ontology analysis of upregulated genes in dT0-3_failed TLSs. The top 15 ontologies were selected. **d,** Venn diagram showing the overlap of upregulated genes in dT0-3 TLSs and Cluster 8 genes. **e,** Venn diagram displaying the overlap of upregulated genes in dT0-3 TLSs and Cluster 5 genes. **f**, Heatmap analysis of Z-score normalized TPM of Cluster 2 and 3 genes in control TLSs and across sample groups.

## Notes

### Competing Interest Statement

The authors have declared no competing interest.

